# Systematic analysis of 1,298 RNA-Seq samples and construction of a comprehensive soybean (*Glycine max*) expression atlas

**DOI:** 10.1101/2019.12.23.886853

**Authors:** Fabricio Brum Machado, Kanhu C. Moharana, Fabricio Almeida-Silva, Rajesh K. Gazara, Francisnei Pedrosa-Silva, Fernanda S. Coelho, Clícia Grativol, Thiago M. Venancio

**Author notes:** These authors contributed equally to this work (Fabricio Brum Machado, Kanhu C. Moharana). Corresponding authors Av. Alberto Lamego 2000 / P5 / 217; Parque Califórnia Campos dos Goytacazes, RJ Brazil CEP: 28013-602 TMV; KCM.

## Abstract

Soybean (*Glycine max* [L.] Merr.) is a major crop in animal feed and human nutrition, mainly for its rich protein and oil contents. The remarkable rise in soybean transcriptome studies over the past five years generated an enormous amount of RNA-seq data, encompassing various tissues, developmental conditions, and genotypes. In this study, we have collected data from 1,298 publicly available soybean transcriptome samples, processed the raw sequencing reads, and mapped them to the soybean reference genome in a systematic fashion. We found that 94% of the annotated genes (52,737/56,044) had detectable expression in at least one sample. Unsupervised clustering revealed three major groups, comprising samples from aerial, underground, and seed/seed-related parts. We found 452 genes with uniform and constant expression levels, supporting their roles as housekeeping genes. On the other hand, 1,349 genes showed heavily biased expression patterns towards particular tissues. A transcript-level analysis revealed that 95% (70,963/74,490) of the known transcripts overlap with those reported here, whereas 3,256 assembled transcripts represent potentially novel splicing isoforms. The dataset compiled here constitute a new resource for the community, which can be downloaded or accessed through a user-friendly web interface at http://venanciogroup.uenf.br/resources/. This comprehensive transcriptome atlas will likely accelerate research on soybean genetics and genomics.

## Introduction

Soybean (*Glycine max* [L.] Merr.) is one of the most important legume crops worldwide. It is critically important in human nutrition, animal feed, and biotechnological applications. Global climate change and increased food demand resulting from a growing human population have been fueling the development and application of biotechnological methods to generate better cultivars (Iizumi et al., 2014). In recent years, various omics approaches have been deployed to improve productivity of several crops, including soybean. An important achievement in soybean omics-based research was the availability of whole-genome sequencing data, which helped identify molecular markers (e.g. single nucleotide polymorphisms, SNPs) (Schmutz et al., 2010;Deshmukh et al., 2014) that are instrumental in the identification of genes associated with various phenotypes of interest. Further, the soybean whole-genome sequencing project has also contributed to the substantial rise in soybean transcriptome studies (Libault et al., 2010;Severin et al., 2010;Garg and Jain, 2013;O’Rourke et al., 2017), initially dominated by microarray platforms and later by RNA-Seq technologies.

To date, several studies reported spatiotemporal changes occurring in various soybean tissues using RNA-seq. The two first soybean RNA-Seq studies were published by Libault *et al.* (Libault et al., 2010) and Severin *et al.* (Severin et al., 2010). The former reported the sequencing of 14 (mainly root and nodule) tissues, whereas the latter evaluated several tissues and seed developmental stages. Dozens of other studies followed, such as those addressing different life cycle stages (Jones and Vodkin, 2013;Bellieny-Rabelo et al., 2016;Gazara et al., 2019), conditions (Belamkar et al., 2014), and cultivars/lines (Goettel et al., 2014). The accumulation of plant transcriptomic data in public repositories [e.g. Sequence Read Archive (SRA) at the National Center for Biotechnology Information (NCBI)] inspired the development of unified collections or atlases, such as those found for *Arabidopsis thaliana* (Fucile et al., 2011), *Medicago truncatula* (He et al., 2009), *Gl. max* (Supplementary Table S1), as well as multi-species atlases (Dash et al., 2012), which are often reused by the scientific community. Specifically in soybean, Kim *et al.* constructed the SoyNet (www.inetbio.org/soynet) database using 734 microarrays and 290 RNA-seq samples (Kim et al., 2017), while Wu *et al.* uncovered a nodulation-related co-expression module by analyzing 1,270 microarray samples generated with Affymetrix gene chips (Wu et al., 2019).

Despite the previous efforts to integrate soybean transcriptomes, there is a massive amount of soybean RNA-Seq data that remain largely unexplored. Here, we have collected data from 1,298 publicly available soybean RNA-seq samples from the NCBI SRA database. We systematically processed and mapped sequencing reads to the soybean reference genome. Transcriptional levels were estimated to allow a systematic global gene expression analysis, aiming to elucidate the dynamics of transcriptional regulation across this broad range of samples, tissues, and cultivars. Further, the collected and processed data are readily available to allow both, automatic analysis and single-gene investigations using an easy-to-use interface at our lab website (http://venanciogroup.uenf.br/resources/).

## RESULTS AND DISCUSSION

### Data gathering, processing, and mapping to the reference genome reveal an overall high quality of the publicly available soybean RNA-Seq data

We performed an extensive literature mining process to gather as many as possible soybean RNA-seq datasets. A total of 1,742 raw read sequencing files were downloaded from the NCBI SRA database (Supplementary Table S2). Reads obtained from the same biological sample were combined in a single FASTQ file (or in two files, for paired-end data; *\*_1.fq* and *\*_2.fq*). This resulted in 1,298 samples (65% single-end and 35% paired-end) from 84 BioProjects comprising sixteen different broad tissue categories in various developmental stages (Supplementary Table S3). Approximately 35% (458/1298) of the samples lacked cultivar/genotype information in SRA. Among the other 840 samples, we found 157 different soybean cultivar names, although this is likely an overestimation because of authors calling the same cultivars with slightly different names during data submission. The cultivar Williams 82, which had the genome sequenced, represented 23% (302/1,298) of the total samples. Leaves were the most abundant tissue, representing 46% (603/1,298) of the samples (Figure 1). Three libraries from unknown tissue sources were excluded. We have also found that 76% (986/1,295) of the libraries were unstranded (Supplementary Table S3).

**Figure 1:**
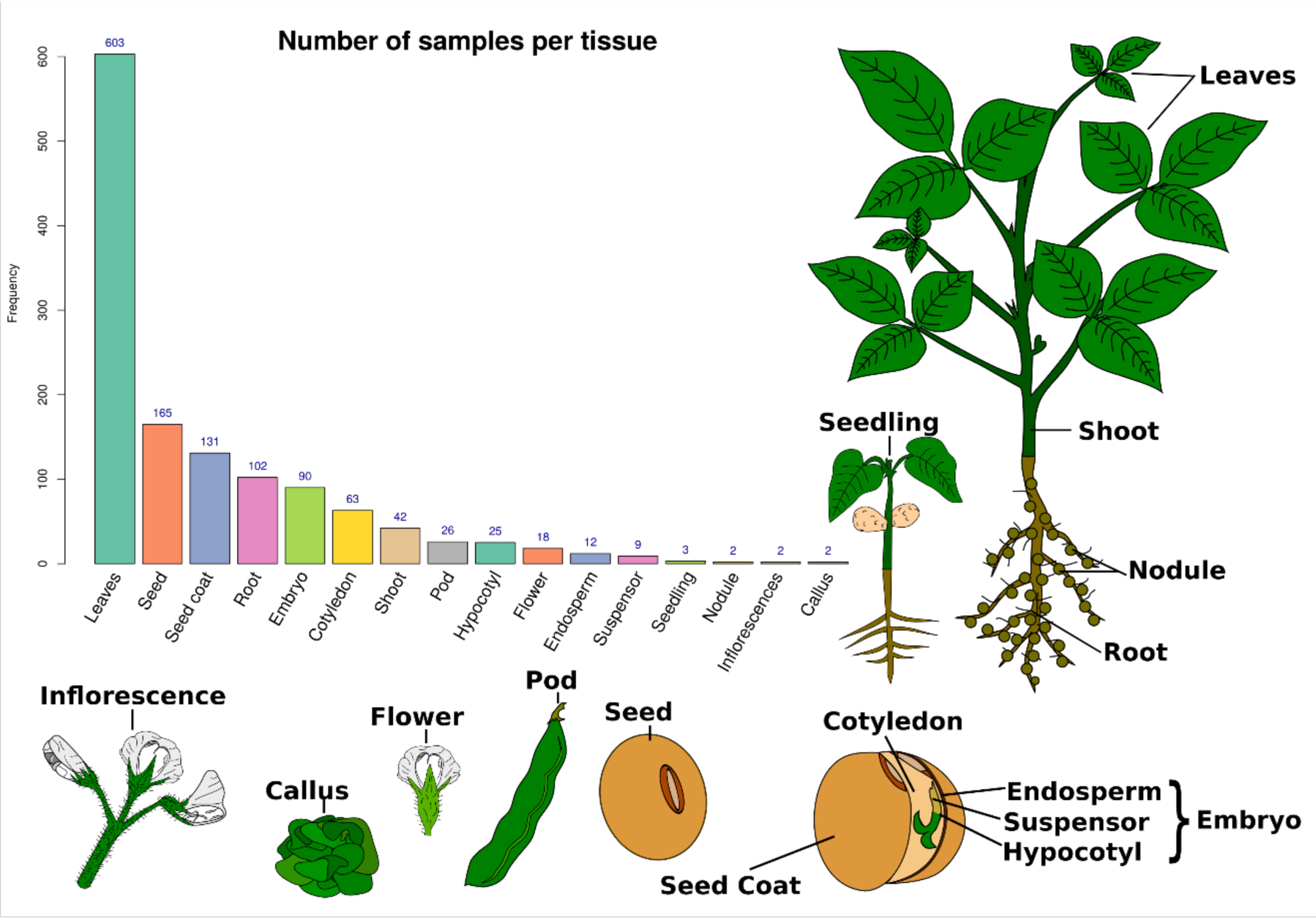
Number of samples analyzed in this study and a graphical representation of each tissue.

Reads from each RNA-seq library were mapped to the reference genome, assembled, and used for estimating gene expression (Figure 2). Whenever present, adapter sequences were trimmed. Reads with average quality lower than 20 were excluded. An average of 32,210,805 million reads pairs per sample with paired-end data and 29,579,316 million reads per sample with single-end data were used for read mapping. Mapped and uniquely mapped reads correspond to an average of 87.9% and 81%, respectively (Supplementary Table S4 and Supplementary Figure 1). Further, we excluded 47 samples for which: i) 50% or more of the reads failed to map or; ii) 40% or more of the reads failed to uniquely map. After these exclusions, 1,248 samples were kept for further downstream analysis.

**Figure 2:**
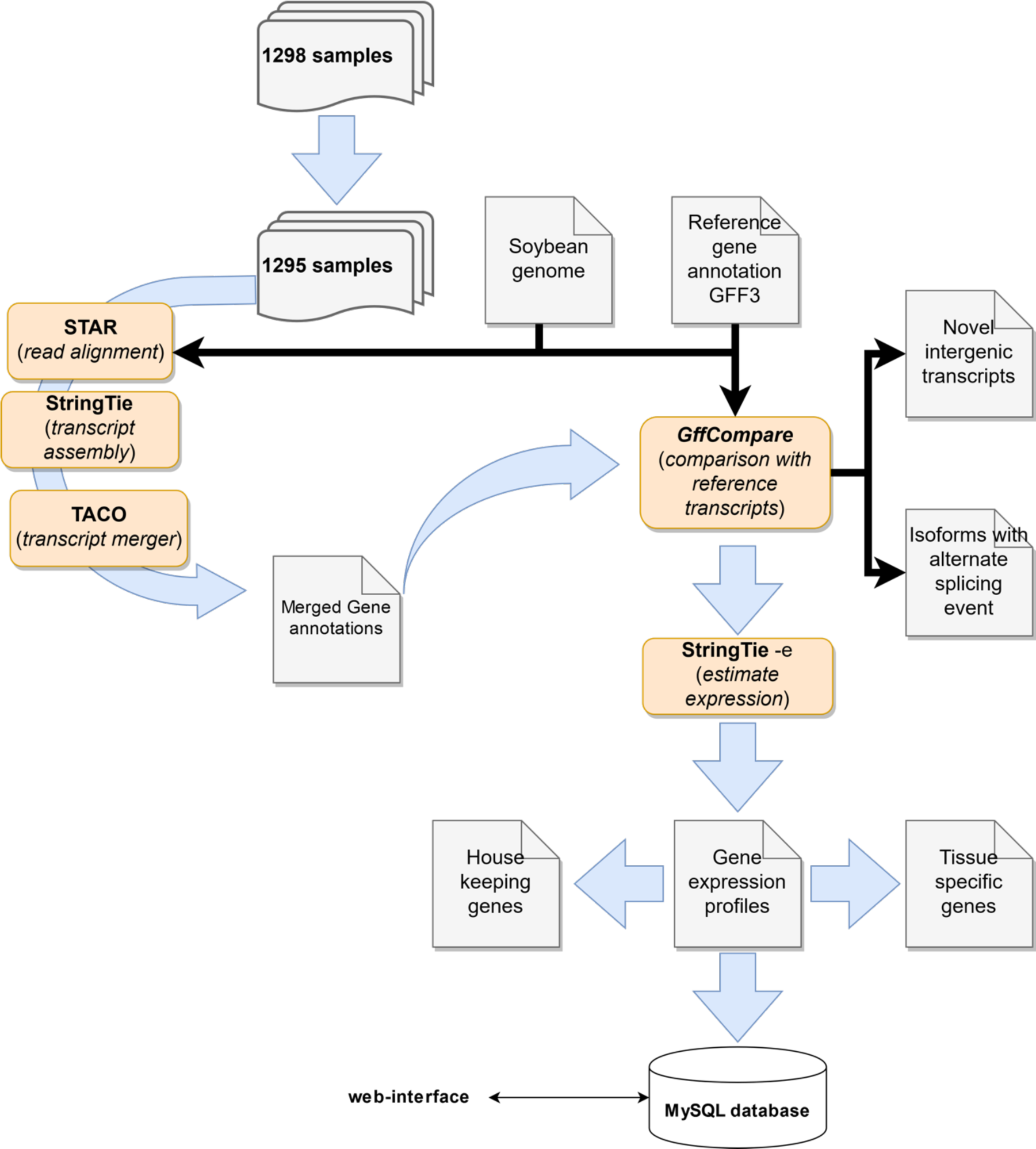
Pipeline used to create the soybean RNA-Seq atlas.

Several methods used to analyze RNA-seq data (e.g. differential gene expression) rely on read count normalization strategies (Robinson and Oshlack, 2010;Po-Yen et al., 2011), such as Reads Per Kilobase Million (RPKM) (Mortazavi et al., 2008), Fragments Per Kilobase Million (FPKM), and Transcripts Per Million (TPM) (Wagner et al., 2012), out of which the latter has been proposed to be more consistent across technical replicates (Wagner et al., 2012;Conesa et al., 2016;Li and Li, 2018). Here, we normalized data using TPM for most of the downstream analysis. Nevertheless, log_2_ transformed raw read counts are more commonly used for quality control steps such as unsupervised sample clustering (Jordan et al., 2015). In addition, many popular tools used for differential gene expression analysis (e.g. DESeq2, edgeR) require raw read counts instead of normalized read counts. Therefore, after read mapping, we estimated transcript abundances in the form of raw read counts per transcript and TPM. Transcript-level expression values were also aggregated to estimate expression at gene level. Gene expression values across 1,248 samples were then used in further downstream analysis.

### Unsupervised sample clustering reveals three major clades comprising underground, aerial, and seed tissues

In transcriptomics studies, gene and samples are often clustered to identify sub-groups with similar transcriptional profiles (Liu and Si, 2014;Marini and Binder, 2019). While gene clustering helps identify co-expressed genes, sample clustering is instrumental to detect broad transcriptional similarities between samples, as well as to identify potential technical artifacts and mislabeled samples. Among several methods, distance-based hierarchical clustering, *K-means* clustering, and dimensional-reduction-based visualization methods (e.g. principal component analysis, PCA) are commonly used. Recently, t-Distributed Stochastic Neighbor Embedding (t-SNE) has been shown to provide a better global structure of sample sub-groups than several other methods (Dey et al., 2017). Here, we employed three sample clustering methods to identify outliers and overall pairwise sample similarity. We used a gene expression matrix as input to perform hierarchical clustering, *K*-means clustering, and t-SNE analysis. These analyses uncovered three major groups comprising samples from aerial, underground, and developmental or seed tissues (Figure 3) (Severin et al., 2010). Interestingly, however, we found an additional cluster comprising samples from leaves and shoots from drought-stress-related and leaf senescence samples. Although not entirely novel, these results are part of an important step to check for technical issues or biases that could, for example, result in the clustering of samples from the same sequencing batch or research group. Four shoot samples and one root sample clustered with seed-embryo samples. After confirming this result with the t-SNE and *K-means* clustering, we excluded these samples. Overall, sample clustering supports a high quality level of the publicly available RNA-Seq samples analyzed here, as only 0.4% (5/1248) of the samples were excluded after the clustering analysis.

**Figure 3:**
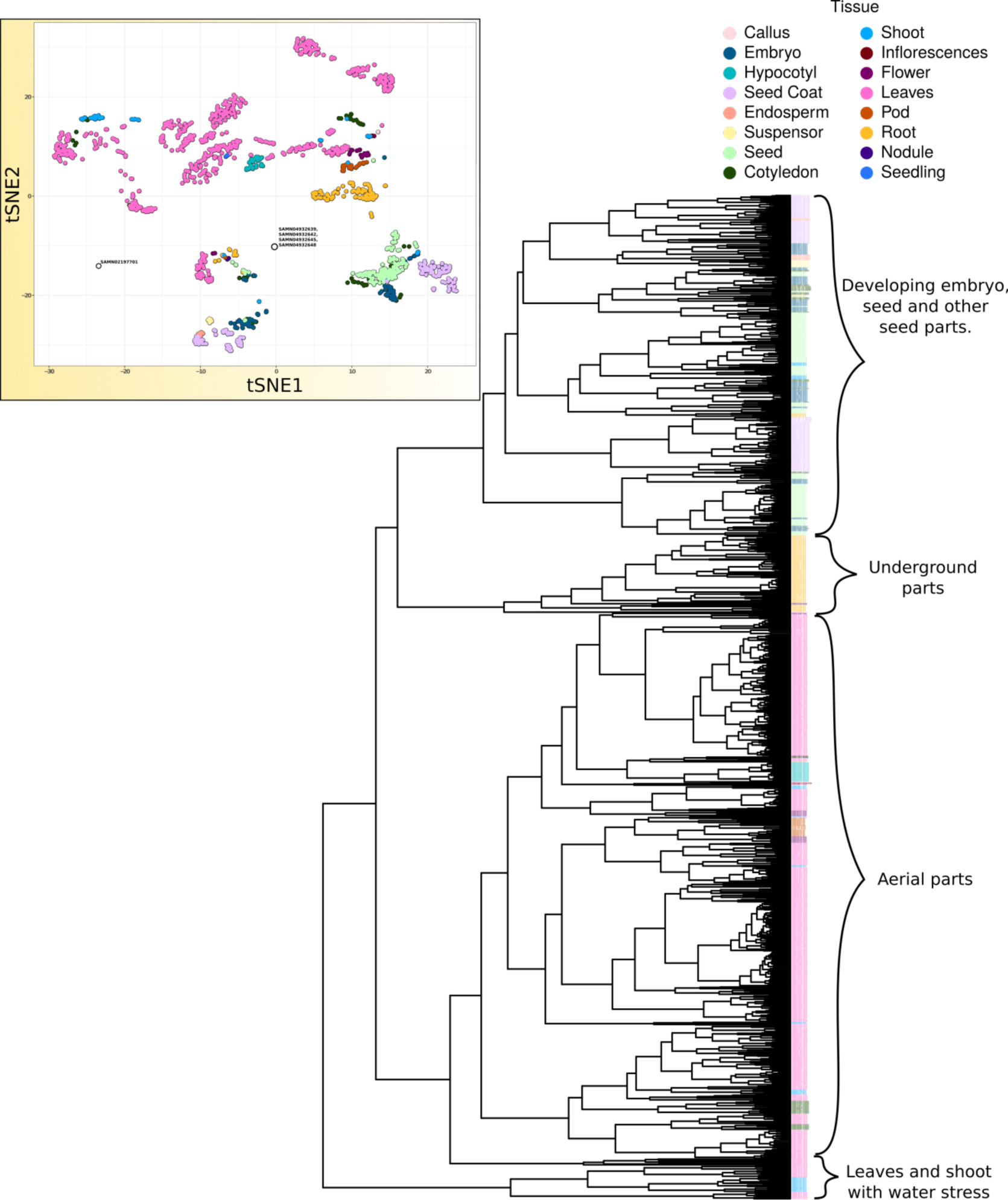
Hierarchical clustering of samples using their transcriptional profiles. Per gene raw read counts were used to perform hierarchical clustering using the R function *hclust()* with default parameters. Samples were grouped into three major clades: aerial, underground, and seed-embryo related. A minor group of samples containing drought-stress-related leaves and shoots was also identified. The upper-left panel shows the sample clustering using t-SNE. Five samples (four from shoot: SAMN04932642, SAMN04932648, SAMN04932639, SAMN04932645 and one from root: SAMN02197701), labeled in the inside plot, showed a very unexpected clustering patterns and were excluded from further analysis. An interactive 3D version of the t-SNE sample clustering is available at http://venanciogroup.uenf.br/resources/.

### Systematic analysis of hundreds of RNA-Seq libraries support the expression of the vast majority of the soybean genes

After comparing the reference transcript annotations (for 56,044 genes) with the merged consensus transcript assembly, we excluded 1.3% (759/56,044) of the genes because of overlapping gene predictions. Next, we applied a minimum TPM threshold of 1 to define a gene as expressed and found that 92.1% (51,644/56,044) of the known soybean protein-coding genes were expressed in at least one sample. The remaining genes had their TPM values set to zero and classified as not expressed. An average of 31,063 genes were expressed per sample. The tissues with the greatest numbers of expressed genes were inflorescence (37,108 genes) and flower (average of 36,051 genes) (Supplementary Figure 2A), whereas nodules had the lowest number of expressed genes (average of 25,718 genes). We also found 16,916 genes expressed in at least 1,150 samples (Supplementary Figure 2B), including 1,758 genes that are expressed in all 1,243 samples. On the other hand, 6% (3,233/56,044) of the genes were not expressed (TPM < 1) in any sample, out of which 82% had coding regions comprising less than 500 codons (Supplementary Figure 3). As a final data quality check, we analyzed the top 1,000 expressed genes from each tissue category using MapMan pathway bins (see Methods). For example, contrasting gene expression profiles of roots and leaves uncovered several expected transcriptional patterns of photosynthesis genes in the latter (Supplementary Figure 4).

**Figure 4:**
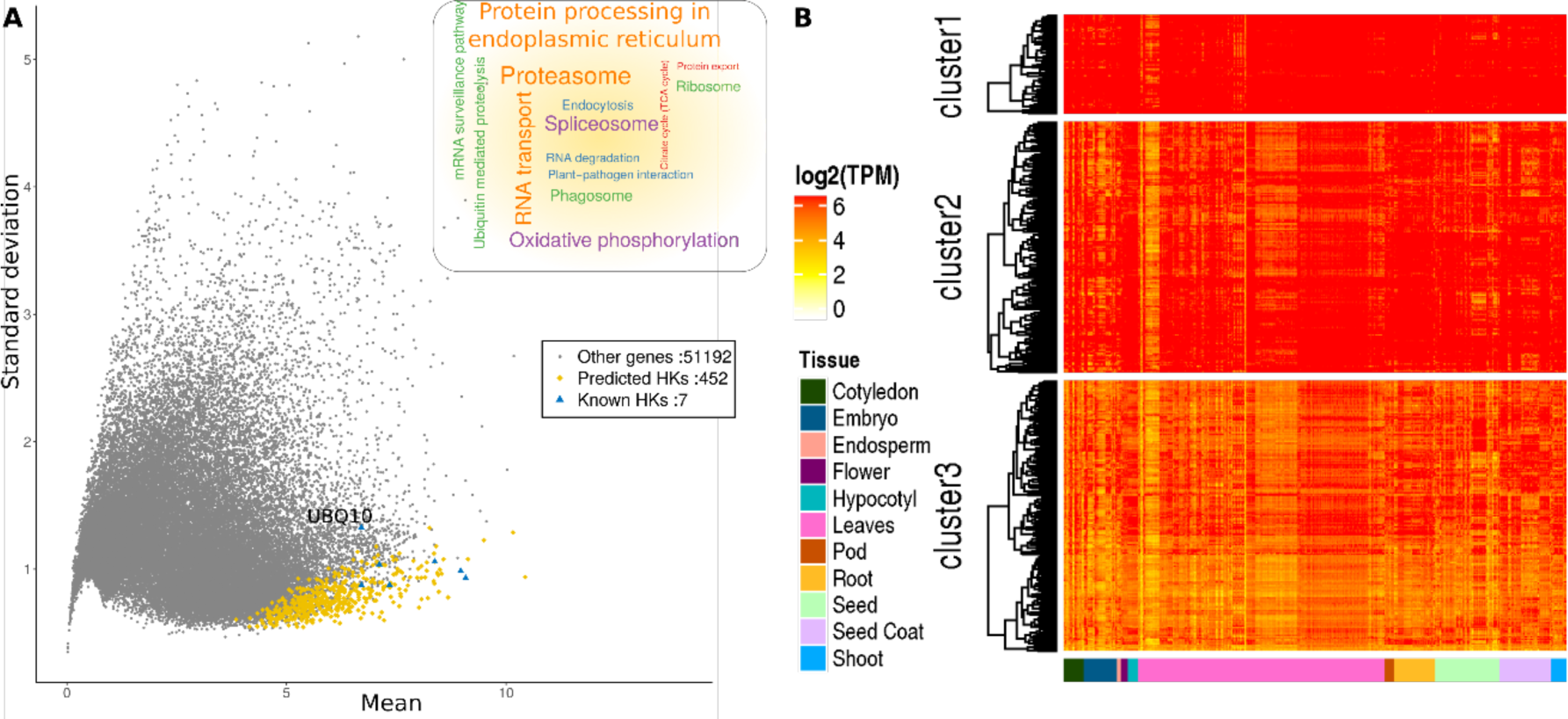
Global gene expression patterns of the housekeeping genes. A. Scatter plot of mean *vs* standard deviation showing uniform and stable expression of 452 housekeeping (HK) genes. The gray dots represent all the non-HK expressed genes (TPM≥ 1 in at least one sample). The word cloud represents KEGG pathways enriched in HK genes (p-value < 0.05). B. Global expression patterns of HK genes. Three main clusters were found with K-means clustering, which were then hierarchically clustered.

### Housekeeping genes

Given the wide coverage of tissues and conditions, we also sought to identify housekeeping (HK) genes based on the assumption that these genes are constitutively and robustly expressed across broad conditions (Czechowski et al., 2005;Hu et al., 2009). Further, several of these genes have also been used as references in real-time quantitative polymerase chain reaction (RT-qPCR) assays (Supplementary Table S5). Hence, by using a large collection of RNA-Seq datasets as the one presented here, one can not only evaluate commonly used reference genes, but also propose new ones. By employing a previously developed method (Hoang et al., 2017), we inferred 452 HK genes (Supplementary Table S6). We evaluated expression levels of each gene in tissues with at least 10 samples and found that HK genes had very low expression variation (Figure 4A). To identify HK genes, we used a score that consists of the product of the Coefficient of Variation and ratio of the maximum to the minimum expression level (see methods for details). Genes with scores within the 1st quartile were classified as HK genes. Further, we used a tissue-specificity index *Tau* (τ) (Yanai et al., 2004;Kryuchkova-Mostacci and Robinson-Rechavi, 2017) to estimate tissue specificity and verify whether our predicted HK genes were broadly expressed or not. The τ values scale from 0 to 1, where low and high values indicate widely expressed and more tissue-specific genes, respectively. The *τ* scores of the HK genes ranged from 0.053 to 0.379, supporting their stable expression level (Figure 6).

**Figure 6:**
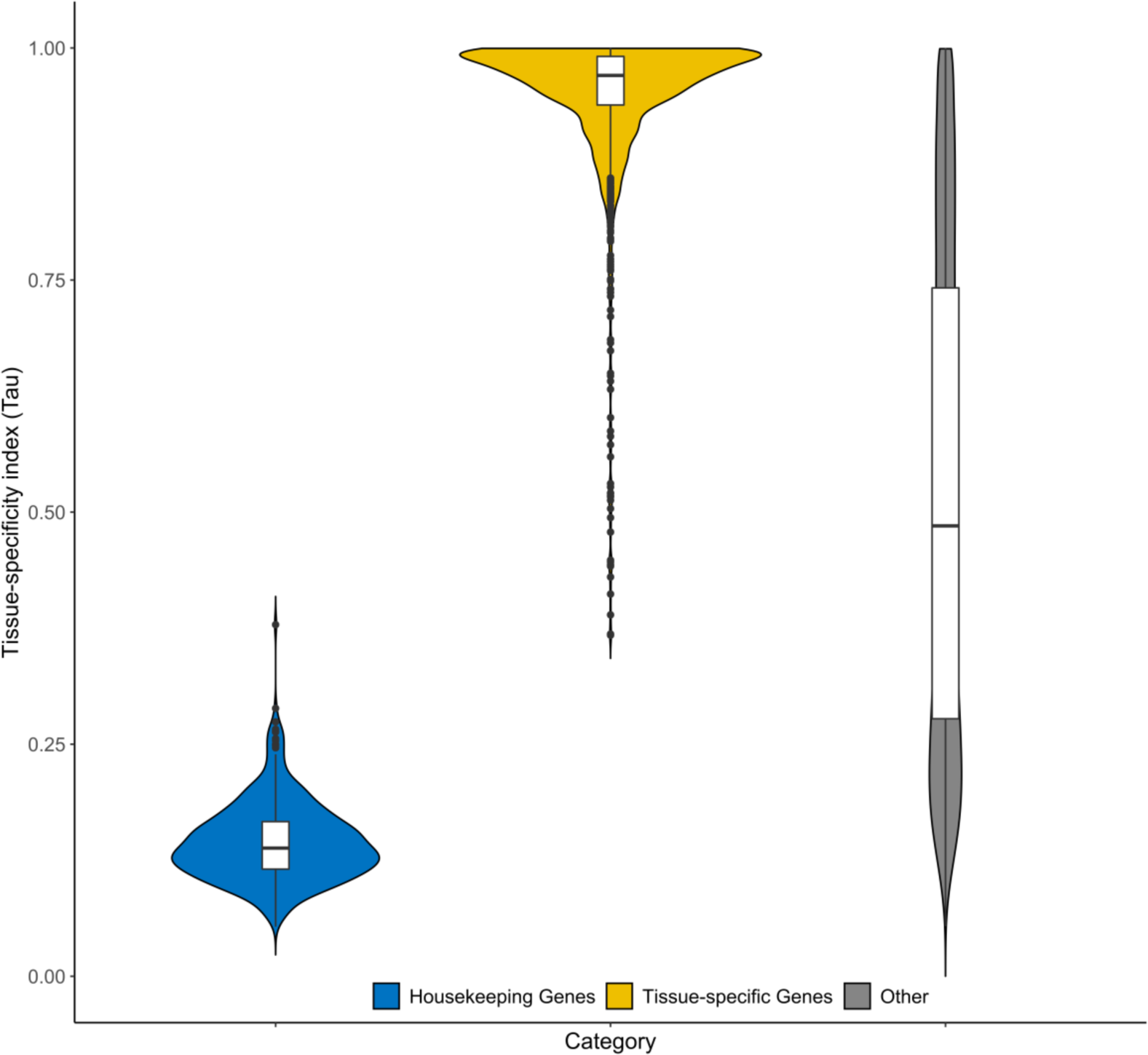
Violin plot showing the distribution of Tau indexes of housekeeping, tissue-specific, and the remaining genes. Tau values range between 0 and 1, with low values indicating a stable and constitutive expression and higher values supporting tissue-specificity.

According to their expression levels, HK genes were grouped in three broad clusters (Figure 4B). Importantly, 7 previously proposed HK genes (Yim et al., 2015) were present in our list (Figure 4), out of which four (*ACT11.C*, *B-actin*, *CYP.B* and*, ELF1α*) belong to cluster 1 (highly expressed, Figure 4A), confirming that high expression is typically an important factor in choosing reference genes. Conversely, given its expression fluctuations (Figure 4), we do not recommend using *UBQ10*, which has also been proposed as a reference gene.

Pathway enrichment analysis of the 452 putative HK genes revealed that these genes are involved in various biological processes such as RNA degradation, mRNA surveillance, and TCA cycle (Figure 4B). We found an enrichment of orthologs of *Arabidopsis* essential genes (Meinke, 2019) among the HK genes (Fisher’s Exact test; p-value = 1.76e-2). Given their roles in basic biological processes, we also verified the conservation of the HK genes in other 14 species on Phytomine and found that 85% (385/452) of them have orthologs in at least 10 other species (Supplementary Table S6), as opposed to an average of 181.6 (± 11.6) in 5 random lists of 452 non-HK genes.

### Tissue-specific gene expression

We compared the global expression patterns between tissues to identify tissue-specific genes (Figure 5). We selected 359 samples that belong to the same tissues and clustered together (Supplementary Table S7), which resulted in the exclusion of four tissue categories. The 12 tissues were compared with each other (a total of 144 comparisons), resulting in a total of 1,349 genes up-regulated in a single tissue as compared to all the others (Figure 7; Supplementary Table S8). Importantly, 96% of these genes (1,300/1,349) had τ indexes greater than 0.8 and median *τ* of 0.9704 (Figure 6). Given their strong preferential expression in particular tissues, we called these genes as tissue-specific.

**Figure 5:**
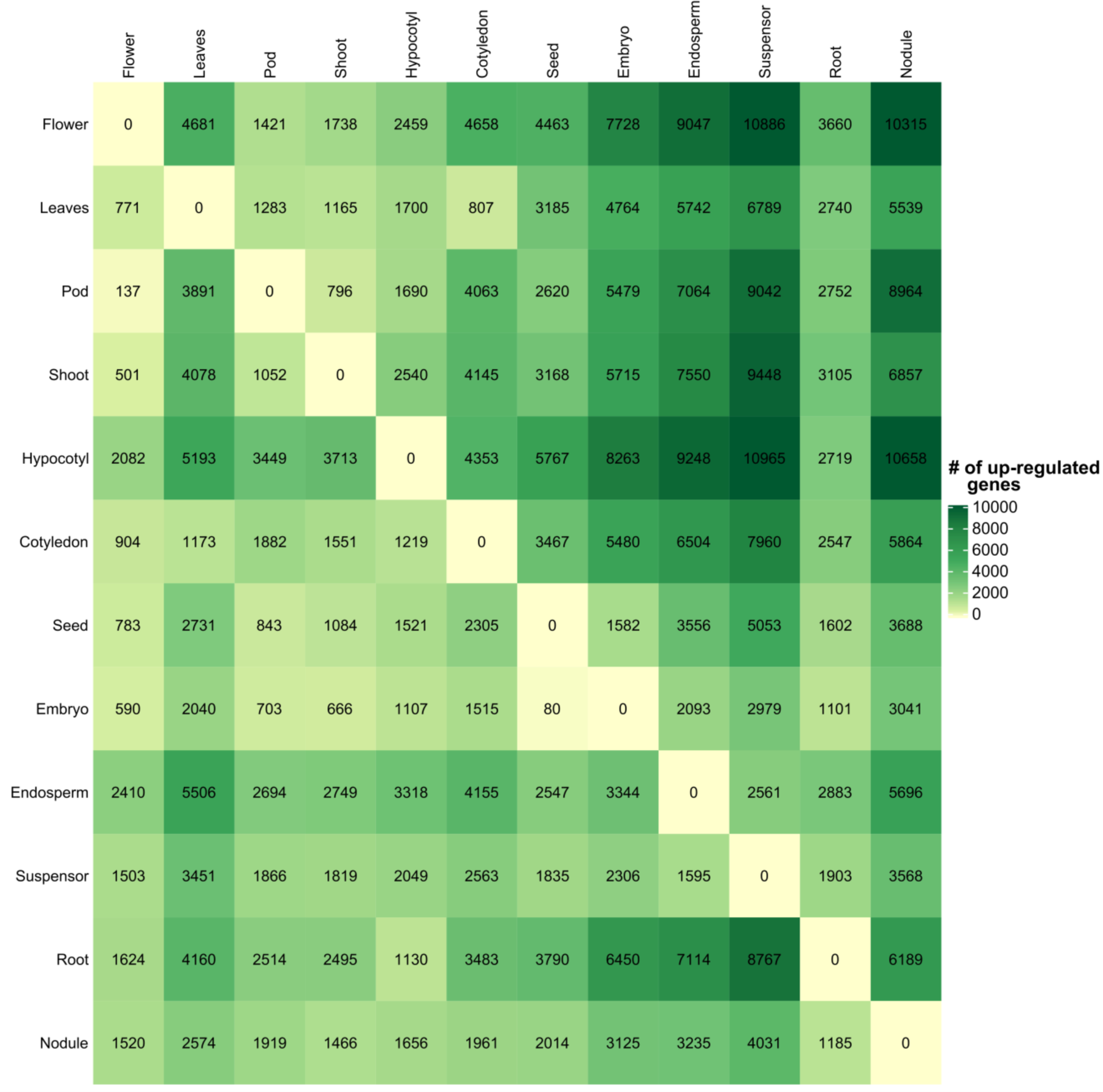
Heatmap showing the number of up-regulated genes in the tissues from the rows when compared with those from the columns. Gene up-regulation was determined by using a log_2_ (fold-change) ≥ 2 and adjusted p-value ≤ 0.05 using the moderated t-statistic in the *limma* package.

**Figure 7:**
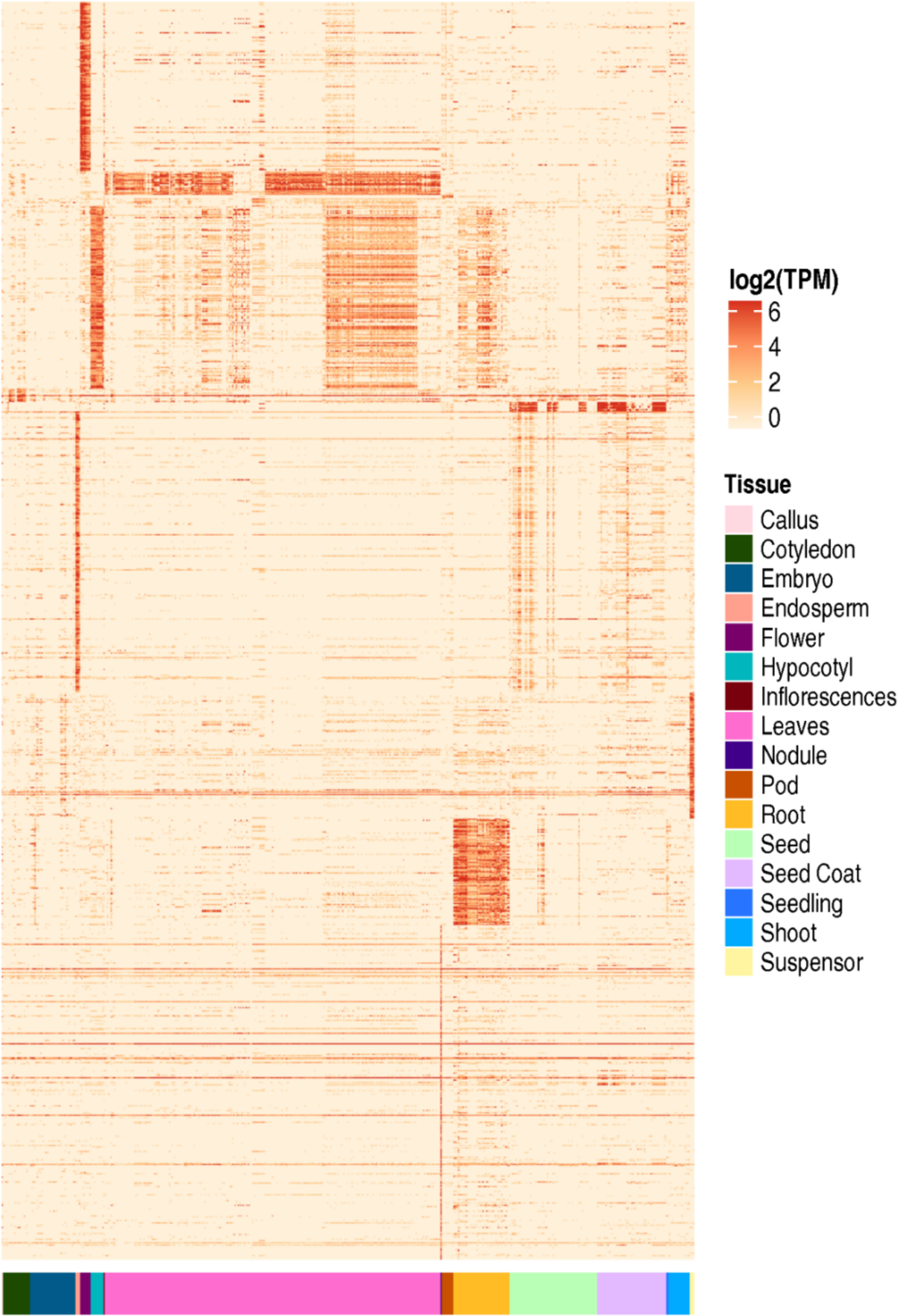
Global transcriptional patterns of tissue-specific genes. Expression values are represented as log_2_(TPM) values in 1243 samples.

The number of tissue-specific genes ranged from 4 in pods to 358 in nodules. Collectively, nodule (26.5%) and endosperm (301; 22%) account for nearly half of the tissue-specific genes. The lower number of tissue-specific genes in leaf, shoot, cotyledon, and pod can be explained by the physiological or developmental relatedness of some samples (e.g. cotyledon and seed). Notably, 39% (520/1,349) of the tissue-specific genes identified here were also identified by Severin et. al (Severin et al., 2010) using a much smaller set of samples, supporting the general high quality and reproducibility of the publicly available soybean transcriptomes. Strikingly, nearly 12% (168/1,349) of the tissue-specific genes were transcription factors (TFs) (Table 1), which is a remarkable enrichment (Fisher’s Exact Test, p-value = 2.94e-11) considering the overall abundance of TFs in the soybean genome (Moharana and Venancio, 2019). Among the tissue-specific TFs, 27, 21, and 20 genes belong to the MYB, C2H2, and ERF families, respectively. Of the 27 MYB TFs, 20 were specific to flower (n=8), hypocotyl (n=7), and endosperm (n=5). Of the 21 C2H2 genes, 12 were specific to nodule (n=6) and endosperm (n=6). Ten out of 20 ERF genes and six out of 10 WRKY genes were specific to hypocotyl. Finally, 8 of 9 MIKC type MADS TFs were flower-specific. Several interesting tissue-specific genes are discussed in the sections below.

**Table 1:** Tissue-specific transcription factors.

| Transcription factor family | Cotyledon | Endosperm | Flower | Hypocotyl | Leaves | Nodule | Pod | Root | Seed | Shoot | Suspensor | Total |
| --- | --- | --- | --- | --- | --- | --- | --- | --- | --- | --- | --- | --- |
| MYB |  | 5 | 8 | 7 | 2 |  | 1 | 2 | 1 |  | 1 | 27 |
| ERF |  | 1 | 1 | 10 |  | 3 |  | 3 |  |  | 2 | 20 |
| C2H2 |  | 6 |  | 1 |  | 6 | 2 | 2 |  |  | 4 | 21 |
| NAC |  |  |  | 2 |  | 1 |  |  | 1 |  | 4 | 8 |
| bHLH | 2 | 1 |  | 2 |  |  |  | 4 |  |  |  | 9 |
| WRKY |  |  |  | 6 |  |  |  | 2 |  |  | 2 | 10 |
| MYB_related |  | 2 | 1 | 1 |  |  |  |  |  |  |  | 4 |
| LBD |  |  | 1 |  |  |  |  | 1 |  |  | 1 | 3 |
| G2-like | 1 | 1 |  |  |  |  |  | 1 |  |  |  | 3 |
| NF-YB |  | 1 |  |  |  | 2 |  |  |  |  |  | 3 |
| M-type |  | 2 |  |  |  | 1 |  |  |  |  |  | 3 |
| MIKC |  |  | 8 |  |  |  |  | 1 |  |  |  | 9 |
| HD-ZIP |  | 2 |  |  |  |  |  |  |  |  | 2 | 4 |
| GRAS |  |  |  | 1 |  | 2 |  |  |  |  |  | 3 |
| bZIP |  | 2 |  |  |  | 4 |  |  |  |  |  | 6 |
| B3 |  | 2 |  |  |  |  |  |  |  |  | 2 | 4 |
| AP2 |  |  |  |  |  | 2 |  |  |  |  | 1 | 3 |
| ZF-HD |  | 2 |  |  |  |  |  |  |  |  |  | 2 |
| YABBY |  |  | 1 |  |  |  |  |  |  |  |  | 1 |
| WOX |  |  |  |  |  |  |  |  |  |  | 3 | 3 |
| SRS |  |  |  |  |  | 1 |  |  |  |  |  | 1 |
| SBP |  |  |  |  |  |  |  |  |  | 1 |  | 1 |
| NZZ/SPL |  | 2 |  |  |  |  |  |  |  |  |  | 2 |
| Nin-like |  |  |  |  |  | 6 |  |  |  |  |  | 6 |
| NF-YC |  | 3 |  |  |  |  |  |  |  |  |  | 3 |
| NF-YA |  |  |  |  |  | 1 |  |  |  |  |  | 1 |
| HSF |  |  |  | 1 |  |  |  |  |  |  |  | 1 |
| GRF |  |  |  |  |  |  |  |  |  | 1 |  | 1 |
| GATA | 1 |  |  |  |  |  |  |  |  |  |  | 1 |
| Dof |  |  |  | 1 |  |  |  |  |  |  |  | 1 |
| CPP |  | 1 |  |  |  |  |  |  |  |  |  | 1 |
| C3H |  | 3 |  |  |  |  |  |  |  |  |  | 3 |
| **Total** | **4** | **36** | **20** | **32** | **2** | **29** | **3** | **16** | **2** | **2** | **22** | **168** |

### Nodule-specific genes

Symbiotic N_2_ fixation takes place in root nodules of several Fabaceae species. Nodulation had a single origin in the common ancestor of the N_2_-fixing clade, followed by multiple independent losses (Griesmann et al., 2018). Among the genes lost in non-nodulating species, *Nodule Inception* (NIN) and *Rhizobium-Directed Polar Growth* (RPG) were reported to be of paramount importance for the origin of root nodules (Griesmann et al., 2018). As mentioned above, nodule is the tissue with the greatest number of tissue-specific genes in soybean, a trend that has also been reported in other legumes (Benedito et al., 2008). Soybean nodules have been shown to correlate poorly with other tissues at the transcriptional level (Severin et al., 2010), a finding that we corroborated here.

We found several nitrogen fixation genes as nodule-specific, including two leghemoglobin (*Glyma.10G199000, Glyma.20G191200*) and ten nodulin genes. The TF families mostly represented among the 29 nodule-specific TFs were NIN-like (n=6) and C2H2 (n=6). A higher percentage of NIN-like and C2H2 nodule-specific TFs have been also described previously (Libault et al., 2010;Severin et al., 2010). Importantly, NIN-like and C2H2 TFs are important in nitrate signaling (Konishi and Yanagisawa, 2013) and symbiosome differentiation during nodule development (Sinharoy et al., 2013). We also found three nodule-specific ERF TFs that are conserved in *Phaseolus vulgaris* and *Medicago truncatula* and are essential for nodule differentiation and development (Vernié et al., 2008).

We found 12 soybean nodule-specific genes within the experimentally validated list of over 200 nodulins described previously (Roy et al., 2019). These 12 genes include the above mentioned ERF TFs, NIN (*Glyma.04G000600*), C2H2 (*Glyma.07G135800*), and GRAS (*Glyma.16G008200*). Next, we analyzed the 28 genes from a nodule-related module identified in a co-expression network derived from soybean microarray data (Wu et al., 2019). Notably, 9 of these 28 genes were identified as nodule-specific in our analysis: one leghemoglobin (*Glyma.10G199000*), two NIN-like TFs (*Glyma.02G311000, Glyma.14G001600*), two purine biosynthesis genes (*Glyma.08G001000, Glyma.11G221100*), one iron transporter (*Glyma.05G121600*), one zinc finger protein-related (*Glyma.08G044700*), one sulfate transporter (*Glyma.18G018900*), and a formyl transferase (*Glyma.19G115900*).

### Endosperm-specific genes

The endosperm plays important roles during seed development. *Ar. thaliana* endosperm-specific genes are associated with cell cycle, DNA processing, chromatin assembly, protein synthesis, cytoskeleton-and microtubule-related processes, and cell/organelle biogenesis and organization (Day et al., 2008). Out of the 301 endosperm-specific genes reported here, 9 (*Glyma.19G040600, Glyma.09G194500, Glyma.01G147300, Glyma.19G058100, Glyma.19G044000, Glyma.04G187100, Glyma.03G219800, Glyma.02G255900*, and *Glyma.08G129200*) encode chromatin modifiers such as histone acetyltransferases, histone-lysine n-methyltransferases, histone deacetylases, and histone demethylases. Further, 17 endosperm-specific genes encode F-box proteins and 8 genes encode BTB-POZ and MATH domain proteins, which likely operate in the ubiquitin-proteasome pathway (Smalle and Vierstra, 2004;Figueroa, 2005). We also found 36 endosperm-specific TFs, including 6 and 5 C2H2 and MYB TFs, respectively. Together, these results clearly show a number of endosperm-specific genes as involved in transcriptional and post-transcriptional regulatory processes.

### Flower-specific genes

The genetic basis of floral development has been widely studied in several plants, including *Ar. thaliana* and *Antirrhinum majus* (Soltis et al., 2007;Bowman et al., 2012). According to the ABCDE model, most of the genes involved in the regulation of flower development encode MADS and AP2/ERF TFs (Chi et al., 2017). The combinatory action of these genes regulates the development of various distinct floral parts. For example, *Ar. thaliana* sepal development is regulated by the MADS-box gene *APETALA1* (AP1) together with the ERF TF *APETALA2* (AP2). Similarly, two MADS-box genes, *APETALA3* (AP3) and *PISTILLATA* (PI), regulate petal/stamen development, whereas the MADS-box gene *AGAMOUS* (AG) regulates carpel development. These basic regulators of flower development are also conserved in other angiosperms (Becker, 2003;Zhao et al., 2017). Further, 491 genes have been suggested to be involved in soybean flower development (Jung et al., 2012).

Recently, several studies reported transcriptional changes during flowering time in legumes (Weller and Ortega, 2015). We found 182 flower-specific genes, including at least 20 members of the *plant invertase/pectin methylesterase inhibitor* (PMEI) superfamily, which is involved in cell wall modification in *Ar. thaliana* (Zhao et al., 2015). Specific PMEIs are highly expressed in specific wheat floral parts, such as anthers and pollen tubes (Rocchi et al., 2012), playing a significant role in flower development (Wormit and Usadel, 2018). In addition, we found 20 flower-specific TFs, mostly from the MYB (40%, 8/20) and MIKC-type MADS (40%, 8/20) families. Finally, out of 8 these MIKC genes, two AGAMOUS-like (*Glyma.03G019400, Glyma.07G081300*) and three PISTILLATA (*Glyma.06G117600, Glyma.13G034100, Glyma.14G155100*) were among the 36 flower-specific genes reported by Jung *et al.* (Jung et al., 2012).

### Identification of novel transcripts

We compared the genomic coordinates of the transcripts assembled in our atlas with those available in Phytozome and categorized them in nine classes (Table 2). We found that 95% (70,963/74,490) of the transcripts precisely matched known transcripts (class =). We also investigated class-J and class-U categories, which account for 3,256 and 23 transcripts, respectively. Class-J comprises multi-exon transcripts with at least one known exon junction, while class-U encompasses transcripts located in intergenic regions. While class-J transcripts include new isoforms of known genes, those from class-U are useful to identify potentially new genes. We found that 30% (983/3256) of the class-J transcripts and 17% (4/23) of the class-U transcripts had TPM ≥ 1 in 907 and 1,207 samples, respectively. Only one of the four class-U expressed transcripts (TU4871, Chr02:12125821-12127123) encode a protein longer than 50 aa, which contains a reverse transcriptase-like RNase_H (PF13456) domain, supporting that it is likely a mobile element. In two of these expressed class-U transcripts (TU28093, TU56508), only one exon showed high read coverage (Supplementary Figure 5).

**Table 2:**
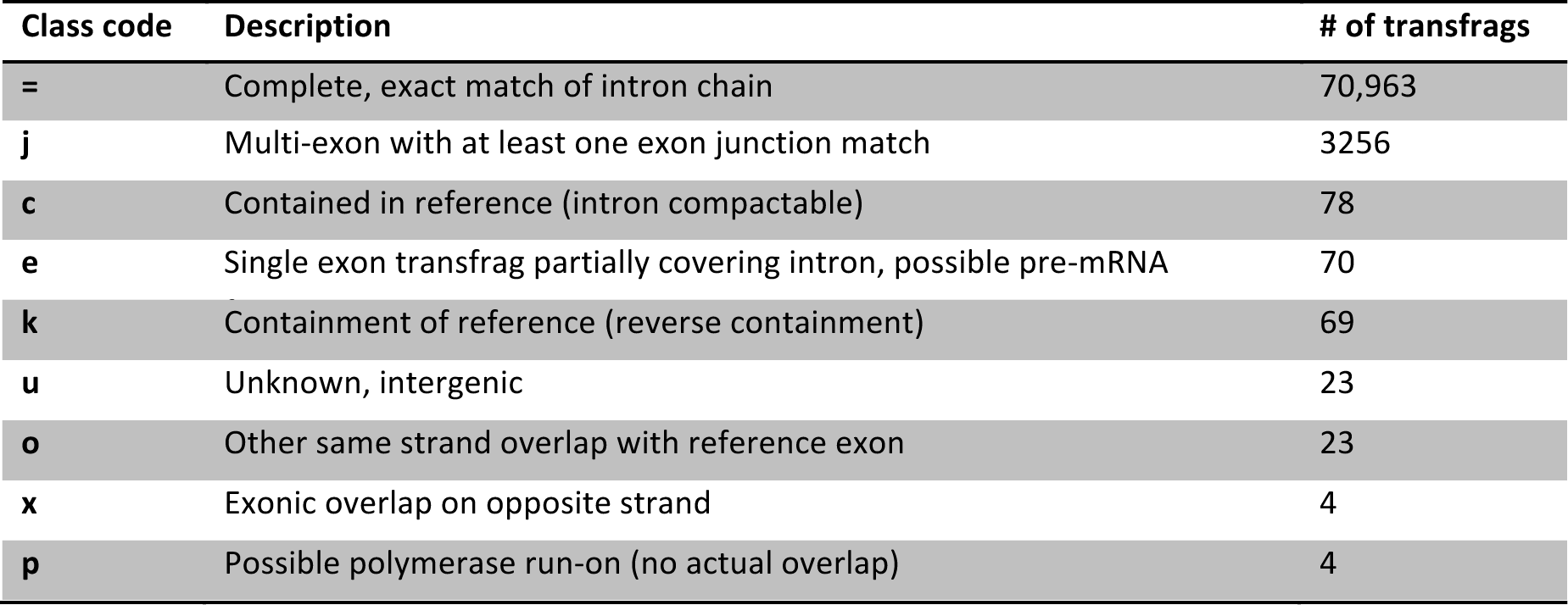
Number of transcripts in each transcript-classification code defined by GffCompare.

All the 3,256 class-J transcripts were further analyzed for alternate splicing (AS) events using ASprofile (Florea et al., 2013). AS events were categorized in one of six categories: (i) exon-skipping; (ii) multiple exon-skipping; (iii) alternative transcription start site (TSS); (iv) alternative transcription termination sites (TTS); (v) intron retention and; (vi) alternate 5’ and/or 3’ exon ends. We detected 6,582 AS events, mostly TSS and TTS (Table 3). Several novel AS events were supported by hundreds of split reads (Supplementary Figure 6-8). For example, TU62356 from *Glyma.17G195900* (CASEIN KINASE 1-LIKE PROTEIN 4) is a novel isoform with a skipped exon (Supplementary Figure 6). Interestingly, we found no support for this alternative isoform in other tissues.

**Table 3:**
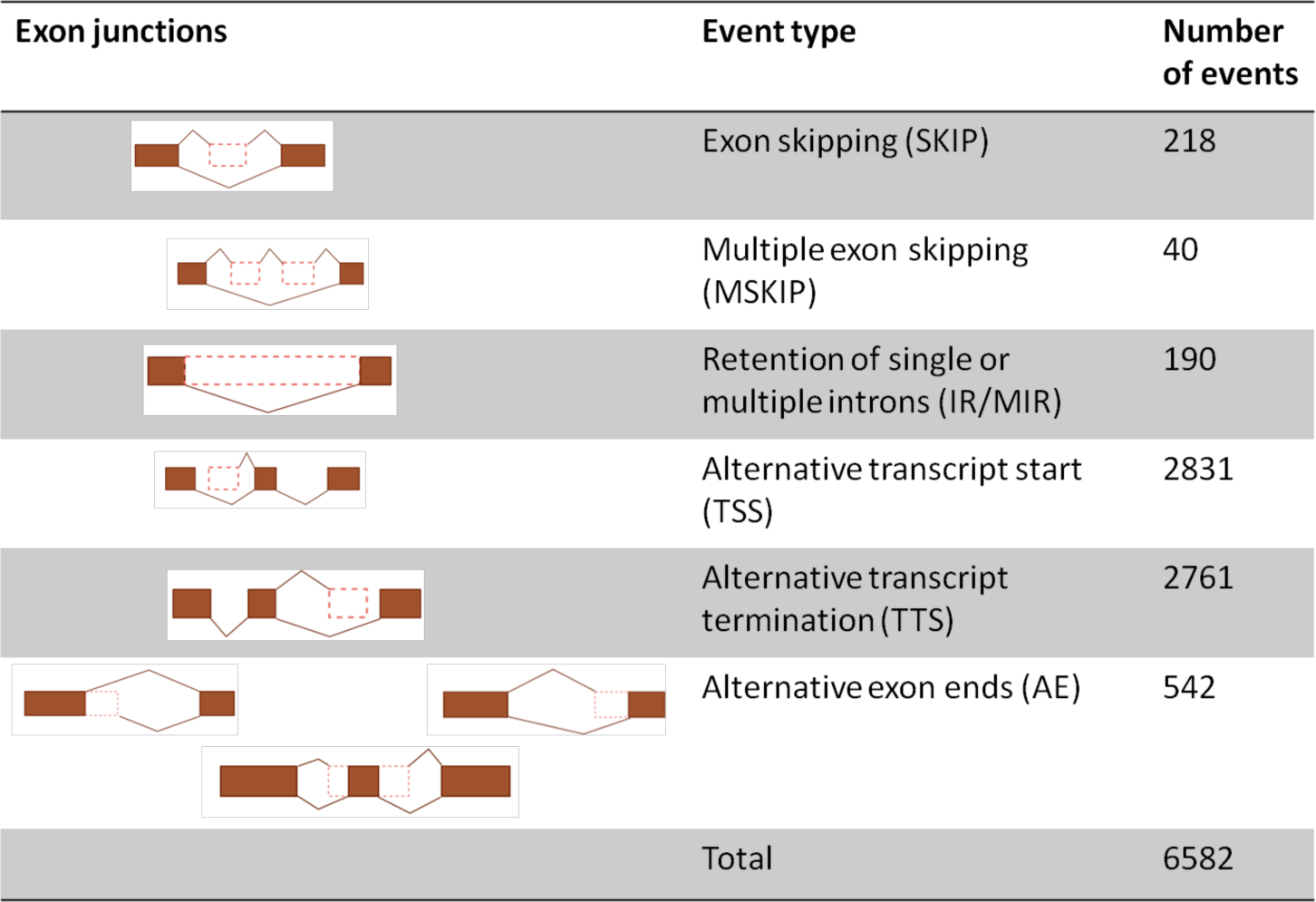
Number of alternative splicing events (AS). The first column illustrates the possible AS isoforms. The boxes represent exons and lines connect adjacent exons in the mature transcript.

| Exon junctions | Event type | Number of events |
| --- | --- | --- |
|  | Exon skipping (SKIP) | 218 |
|  | Multiple exon skipping (MSKIP) | 40 |
|  | Retention of single or multiple introns (IR/MIR) | 190 |
|  | Alternative transcript start (TSS) | 2831 |
|  | Alternative transcript termination (TTS) | 2761 |
|  | Alternative exon ends (AE) | 542 |
|  | Total | 6582 |

### Data availability through a user-friendly web interface

We developed a simple user-friendly web interface to allow researchers to easily explore 1,243 soybean transcriptome samples. Through this interface (Figure 8), one can explore the expression of a particular gene in multiple tissues, with the aid of an image illustrating all the available tissues. Alternatively, users can also retrieve expression profiles of multiple genes in batch, with multiple filtering options (e.g. by tissue, BioProject, study). The outputs can be exported as plain text files. We strongly believe that this website will optimize data reuse and help research groups in their own projects. This service can be freely accessed at http://venanciogroup.uenf.br/resources/.

**Figure 8:**
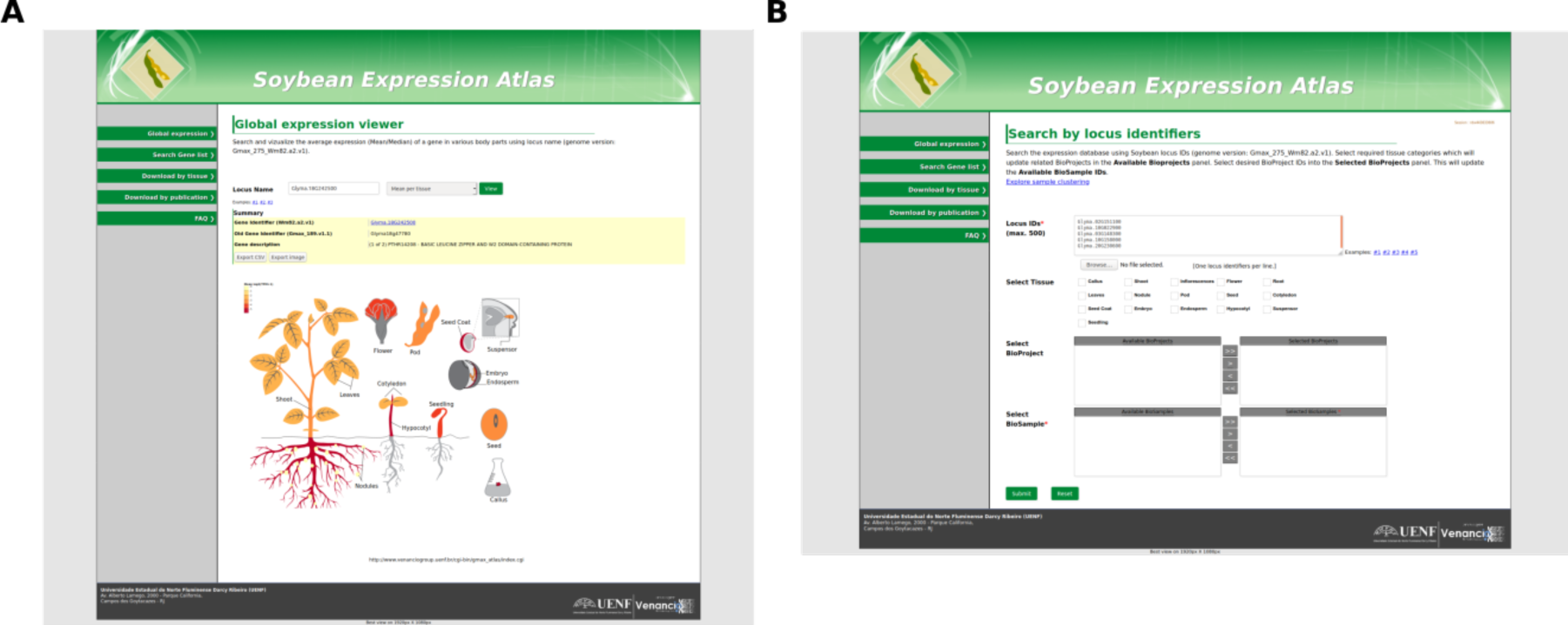
Web interface to browse and download the expression data analyzed in this study. A. Users can search, visualize and download average expression levels in each tissue or; B retrieve expression values in batch in particular samples, tissues, or BioProjects. This resource is available at: http://venanciogroup.uenf.br/resources/.

## Conclusions

We have culled a large collection of publicly available RNA-seq datasets to construct a transcriptome atlas in soybean. We implemented a pipeline with state-of-art methods to map and quantify gene expression levels in 16 different broad tissue categories. This atlas allowed us to identify constitutive and tissue-specific genes. The constitutively expressed genes might, for example, be used as reference genes in RT-qPCR experiments, whereas tissue-specific genes might help scientists test hypotheses in downstream experiments and functional genomics studies. To optimize data reuse, we elaborated a simple web interface to allow the community to quickly access and browse the collected data. We believe this atlas will be an invaluable resource not only for basic research projects, but also in the development of novel strategies to improve soybean productivity to meet increasing global food demands.

## Methods

### Soybean genome and annotation data

Soybean genomic sequences and gene annotation data (assembly version: Gmax_275_Wm82.a2.v1) were obtained from Phytozome (Schmutz et al., 2010;Goodstein et al., 2012). The gene annotation file contained 56,044 and 88,647 genes and transcripts, respectively. The gene annotation file containing exon-intron boundaries (GFF3 format) was used as a reference guide in read mapping. We excluded 759 overlapping genes from the analysis. The gene description file was used to obtain various annotations such as GO, KEGG, KOG, and *Arabidopsis* ortholog descriptions.

### Soybean RNA-Seq data

To identify soybean transcriptome sequencing projects, we searched the NCBI SRA database (https://www.ncbi.nlm.nih.gov/sra) and the metadata were exported by using *Run selector* (https://trace.ncbi.nlm.nih.gov/Traces/study/). We also searched Soybean RNA-seq studies in the literature (up to May 2018) to find additional datasets. We enriched this list of studies with various other details, such as PubMed ID and experiment details obtained by using NCBI *e-fetch*. Using these metadata, we excluded miRNA/siRNA samples and a few other samples showing technical issues such as: i) empty FASTQ files; ii) paired-end samples with single-end reads and; iii) paired-end reads of unequal lengths. Collectively, we downloaded a total of 1,742 *.sra* files (Supplementary table S2), which were decompressed using sra-toolkit (v.2.5.7) (Leinonen et al., 2010).

### Preprocessing and quality control

Quality assessment of FASTQ files was performed using FASTQC (https://www.bioinformatics.babraham.ac.uk/projects/fastqc/). Datasets were processed using Trimmomatic (v0.36) (Bolger et al., 2014) to remove reads with average base quality lower than 20 or containing adapter sequences. Library strandedness was determined with the *infer_experiment.py* script from RSeQC (Wang et al., 2012) using a mapping of 20% of the reads of each sample to the soybean genome in a fast-forward manner using Bowtie2 (Langmead and Salzberg, 2012).

### Transcript assembly and gene expression estimation

We aligned the reads to the *Gl. max* reference genome (Gmax_275_Wm82.a2.v1) by using STAR (v.2.5.3a) (Dobin et al., 2013) with default parameters, along with the soybean gene annotation file containing exon-intron boundaries (in GFF3). When required, STAR also splits reads to find novel exon-intron boundaries or splice sites. The log files were processed to obtain read mapping statistics. Next, StringTie (v. 1.3.4) (Pertea et al., 2015) was used to assemble transcripts and estimate normalized gene expression. We performed transcriptome assemblies for each of the 16 tissues separately. In StringTie, we set the following parameters: i) at least 5 reads with at least 25% of the total read length covering both sides of an exon junction boundary (–j 5 –a 0.25*read_length); ii) average read depth for a transcript of at least 10 (–c 10) and; iii) library strandedness, when applicable. The resulting 16 assembled transcript annotations from each tissue were combined with TACO v0.7.3 (Niknafs et al., 2017). GffCompare (v0.10.5) (https://ccb.jhu.edu/software/stringtie/gffcompare.shtml) was used to compare assembled and reference transcripts. Further, *featureCount* (subread-v1.6.2) (Liao et al., 2014) was used to count the number of reads per feature at transcript and gene levels, while normalized expression was estimated in TPM using StringTie (*–e* option).

### Sample clustering

We assessed the sample clustering patterns by submitting 41,011 genes with mean log_2_ (read count+1) ≥ 1 to: i) hierarchical clustering; ii) t-SNE clustering and; iii) *K-means* clustering. These analyses were performed using R functions (www.r-project.org) *cor*(), *hclust*(), and *kmeans*(). For t-SNE clustering, we used the *t-SNE* R package (Krijthe, 2015) with clustering parameters max_iter= 5000 and perplexity= 50. For hierarchical clustering, sample dissimilarity *(1 – Pearson Correlation Coefficients)* values were used to infer pairwise sample distances. The resulting tree was inspected for unexpected sample clustering patterns. t-SNE separated samples in 35 sub-clusters. Thus, we ran the *K-means* clustering analysis to find 35 centroids (k= 35).

### Identification of novel genes and splicing isoforms

To identify novel genes and isoforms, we analyzed the GffCompare output files. Transcripts not overlapping with any known reference transcript were assigned to class-U. The nucleotide sequences of the class U transcripts were extracted and translated using TransDecoder (v. 3.0.1). Protein domains were predicted using HMMER3 v. 3.1b2 (all default parameters except domain e-value < 0.01) (hmmer.org) and the Pfam database (release 32.0) (El-Gebali et al., 2019). Read coverage of these novel genes were visualized with Gbrowse, available on Soybase (https://soybase.org/gb2/gbrowse/gmax2.0). Class-J transcripts were classified as putative novel isoforms. Splice junctions of these transcripts in GTF format were compared against all known splice junctions using ASprofile v.b-1.0.4 (Florea et al., 2013). The number of reads supporting a splice junction was visualized as sashimi plots using Integrated Genome Viewer (v2.4.10)(Robinson et al., 2011).

### Analysis of the top 1000 highest expressed gene lists

The top 1000 genes with the greatest average TPM in each tissue category were analyzed using MapMan (v3.5.1R2) (Thimm et al., 2004). To assign pathway bins, amino acid sequences of these gene lists were compared against *Arabidopsis* peptide database using Mercator4 (v. 2.0) (Schwacke et al., 2019).

### Identification of housekeeping genes

We selected 11 tissues with at least 10 samples, which resulted in a total of 1,225 samples. The variability in gene expression was evaluated as previously described (Hoang et al., 2017). The following criteria were applied to identify HK genes:

i. A gene with TPM < 1 in a given sample was considered as not expressed (these TPM values were set to 0);
ii. Genes must be expressed in all 1,225 samples. This step resulted in 1,809 genes;
iii. The mean TPM of each gene was calculated by taking the average of the gene expression across all samples;
iv. The Coefficient of Variation (CoV) was computed by taking the standard deviation divided by the mean expression of a gene;
v. The ratio of the maximum to minimum (MFC) was calculated by dividing the largest by the smallest TPM value. A product score (MFC-CoV) was calculated based on the product of CoV and MFC for each gene;
vi. Genes with MFC-CoV scores within the 1^st^ quartile were classified as HK genes.

HK genes were also analyzed using the tissue-specificity index τ (Yanai et al., 2004;Kryuchkova-Mostacci and Robinson-Rechavi, 2017). The τ values ranged from 0 (broad expression) to 1 (exclusive expression). *τ* for each gene was calculated by using the formula:

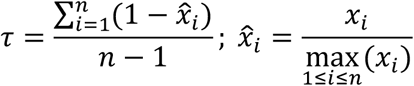

where

*x*_*i*_ = expression of the gene in tissue *i*.

*n* = number of tissues.

### Assessment of tissue-specific expression

We used the log_2_ transformed TPM values for this analysis. Each of the 12 tissues was compared against each other (a total of 144 comparisons) to find significantly over-expressed genes using *limma* (Ritchie et al., 2015). We used log_2_ (fold-change) ≥ 2 and adjusted p-value ≤ 0.05 (moderated t-statistic) to identify significantly over-expressed genes. If a gene *G* is over-expressed in a tissue *T* in comparison to the other 11 tissues, *G* was considered as specifically expressed in *T*. We also used τ to assess tissue-specific expression by applying a minimum threshold of 0.8, as previously recommended (Kryuchkova-Mostacci and Robinson-Rechavi, 2017).

### Gene orthologs and enrichment tests

We obtained the gene descriptions from Phytomine (https://phytozome.jgi.doe.gov/phytomine/begin.do), which is an InterMine (Lyne et al., 2015) interface to genomic data from Phytozome (Goodstein et al., 2012). We used Phytomine to assess the conservation of HK genes in 14 different species (*Ph. vulgaris*, *Me. truncatula, Vigna unguiculata, Ar. thaliana, Oryza sativa, Gossypium raimondii, Carica papaya, Vitis vinifera, Sorghum bicolor, Zea mays, Amborella trichopoda, Selaginella moellendorffii, Physcomitrella. Patens*, and *Volvox carteri*). To estimate the conservation of non-HK genes, we created 5 sets of 452 randomly selected genes from the 55,592 non-HK genes. Each of these sets were searched for orthologs in the above mentioned 14 species. GO enrichment was performed on Phytomine (corrected p-value < 0.05). We performed Kyoto Encyclopedia of Genes and Genomes (KEGG) pathway enrichment using KOBAS 3.0 (Ai and Kong, 2018). We used the Fisher’s Exact test to assess the enrichment of essential genes and TFs in particular gene sets. The list of 510 *Arabidopsis* EMBRYO-DEFECTIVE (EMB) genes (Meinke, 2019) were searched on Phytomine and the corresponding 1,010 soybean orthologs were retrieved. The list of soybean TFs was obtained from a recently published work (Moharana and Venancio, 2019).

### Web server

The TPM and read count values for 54,877 genes across 1243 samples were stored in a relational database implemented in MySQL and hosted on an Apache HTTP web server. The front-end to this database was developed using Python/html/CSS. Interactive visualizations were implemented using *D3.js* (https://d3js.org/) and *Plotly.js* (https://plot.ly/) javascript libraries. The online server is publicly available at http://venanciogroup.uenf.br/resources/.

## Acknowledgements

This work was supported by Fundação Carlos Chagas Filho de Amparo à Pesquisa do Estado do Rio de Janeiro (FAPERJ; grants E-26/010.002019/2014, E-26/102.259/2013, and E-26/203.014/2018), Coordenação de Aperfeiçoamento de Pessoal de Nível Superior - Brasil (CAPES; Finance Code 001), and Conselho Nacional de Desenvolvimento Científico e Tecnológico (CNPq). The funding agencies had no role in the design of the study and collection, analysis, and interpretation of data and in writing.

