## Supplementary figures for "Systematic analysis of 1,298 RNA-Seq samples and construction of a comprehensive soybean (*Glycine max*) expression atlas"

[1] Laboratório de Química e Função de Proteínas e Peptídeos, Centro de Biociências e Biotecnologia, Universidade Estadual do Norte Fluminense Darcy Ribeiro; Campos dos Goytacazes, Brazil.

[#] These authors contributed equally to this work.

[*] Corresponding authors
Av. Alberto Lamego 2000 / P5 / 217; Parque Califórnia
Campos dos Goytacazes, RJ
Brazil
CEP: 28013-602
TMV:; KCM:

### Supplementary figures

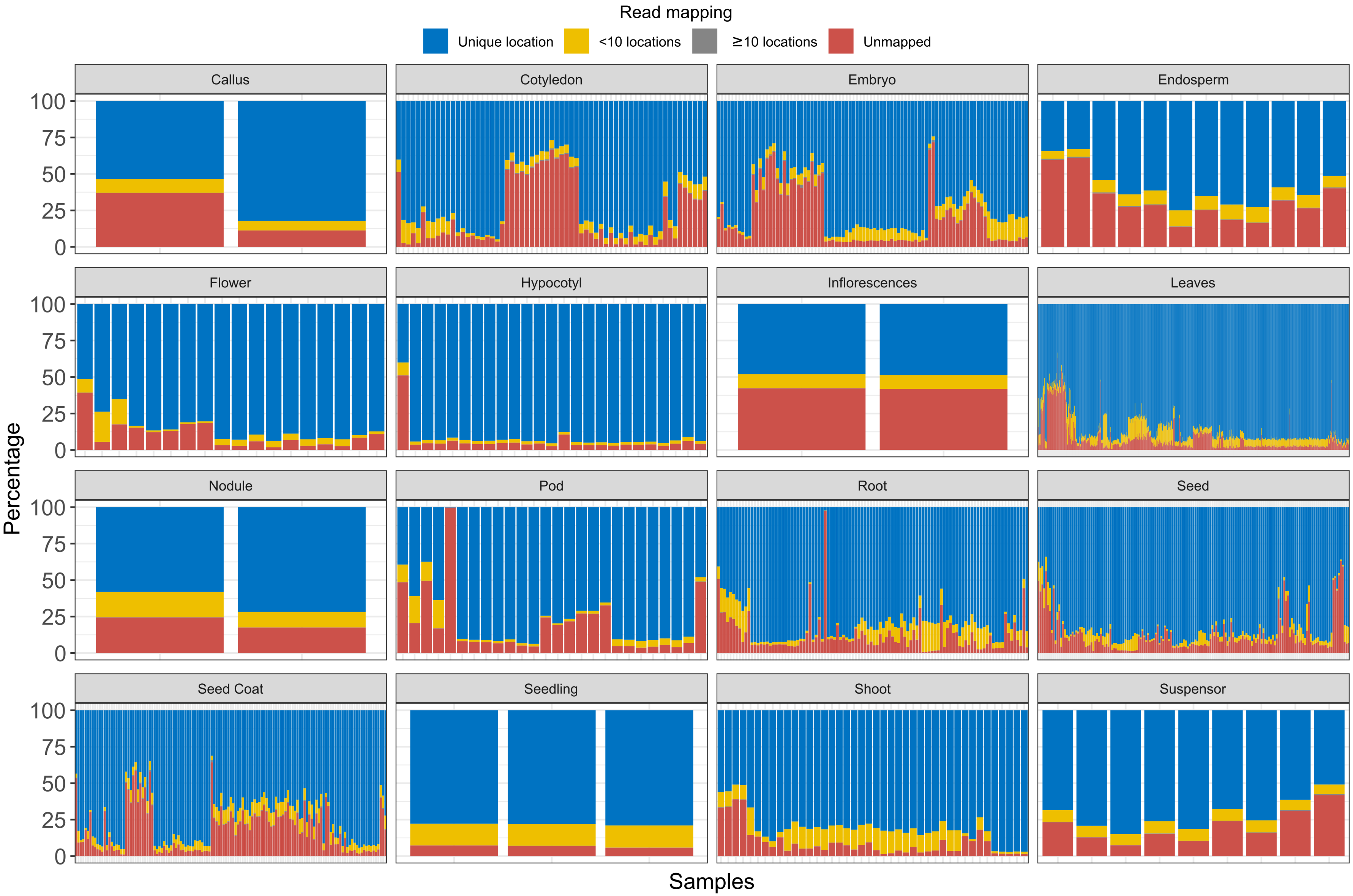


**Supplementary Figure 1:** Stacked histograms showing the read mapping statistics across tissues.

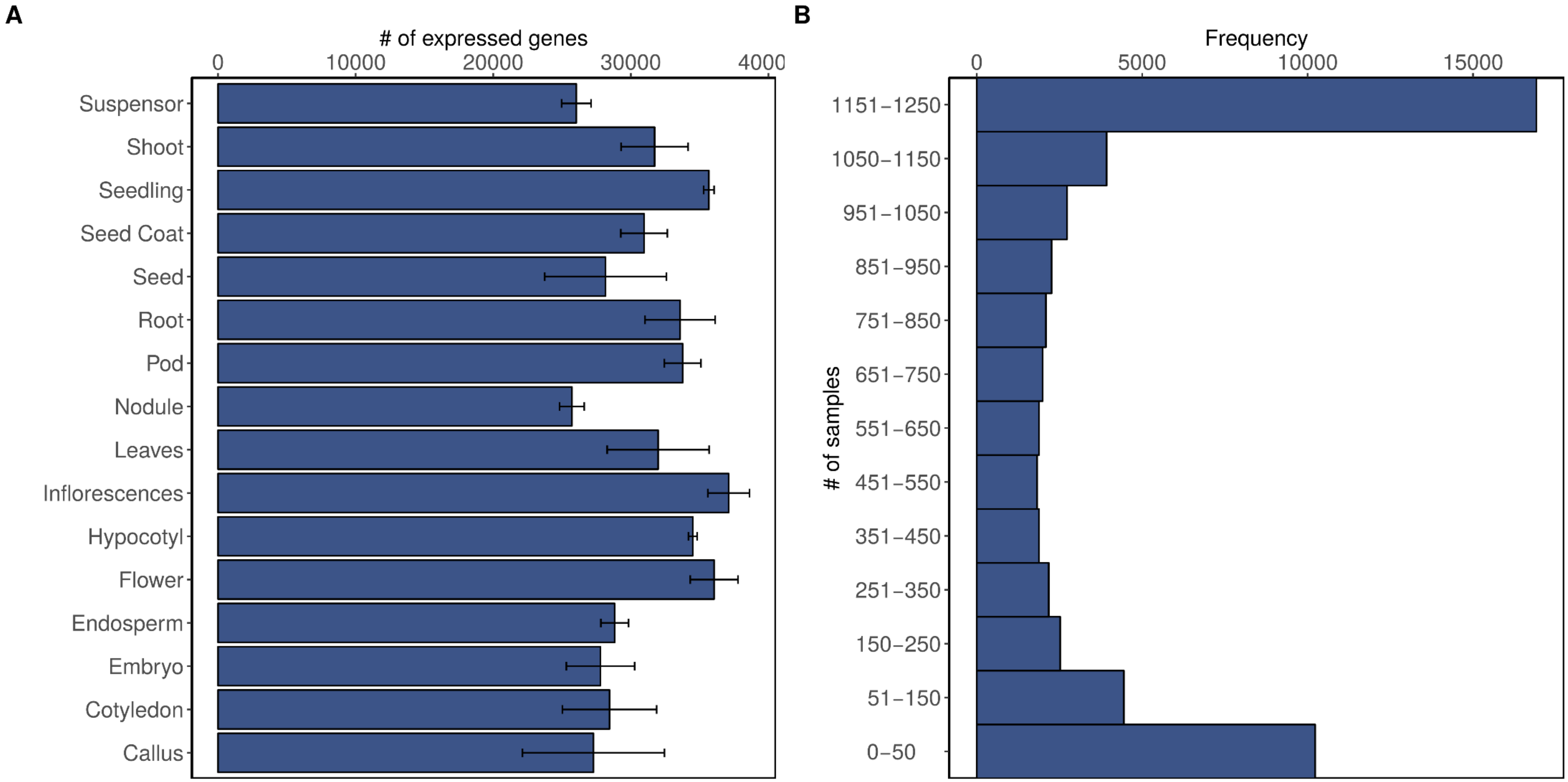


**Supplementary Figure 2:** Tissue- and sample-wise distribution of expressed genes (TPM ≥ 1). A. Number of expressed genes in each tissue. B. Number of samples in which genes are expressed.

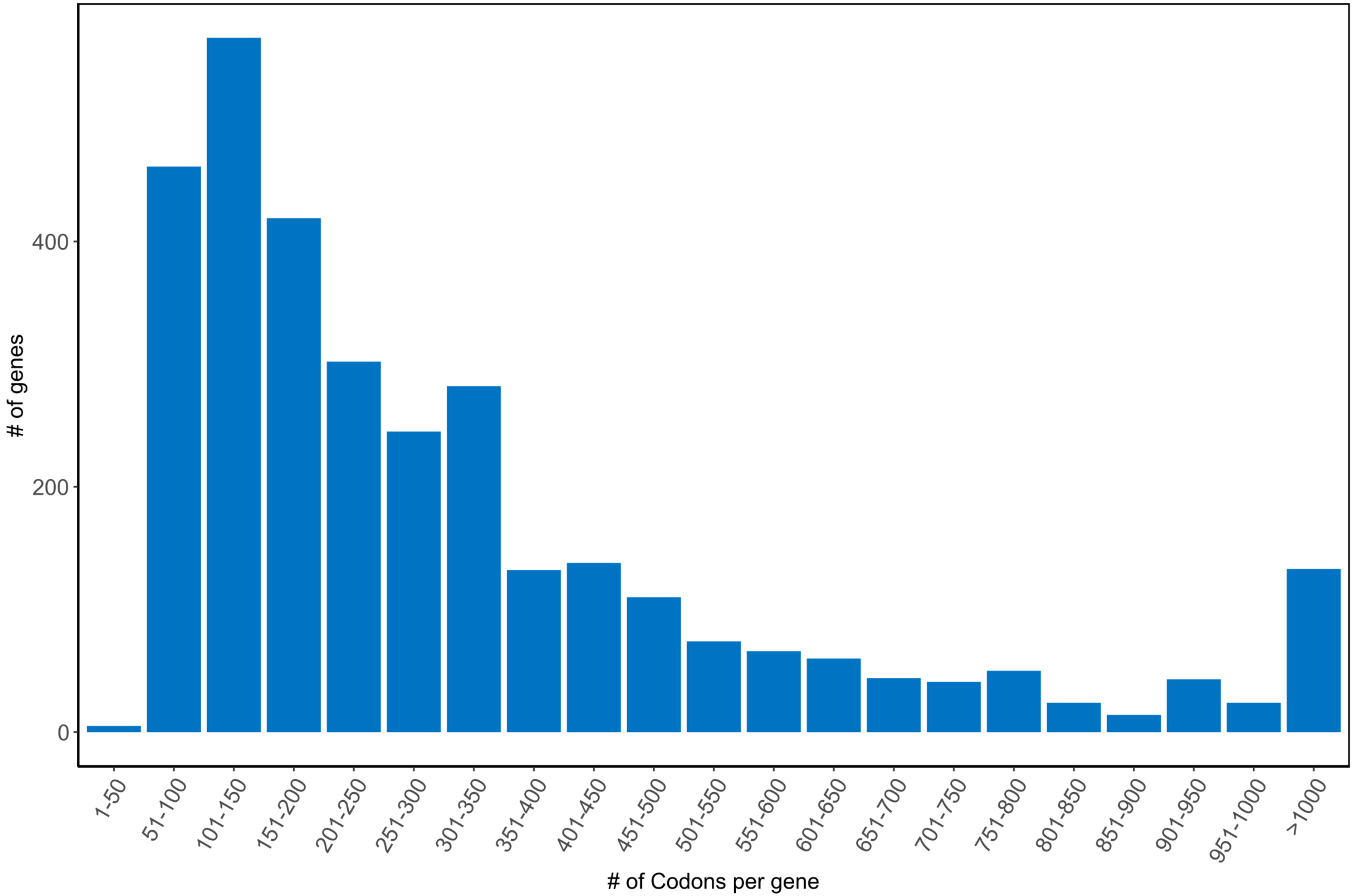


**Supplementary Figure 3:** Length (in codons) of genes with undetectable expression levels (TPM < 1).

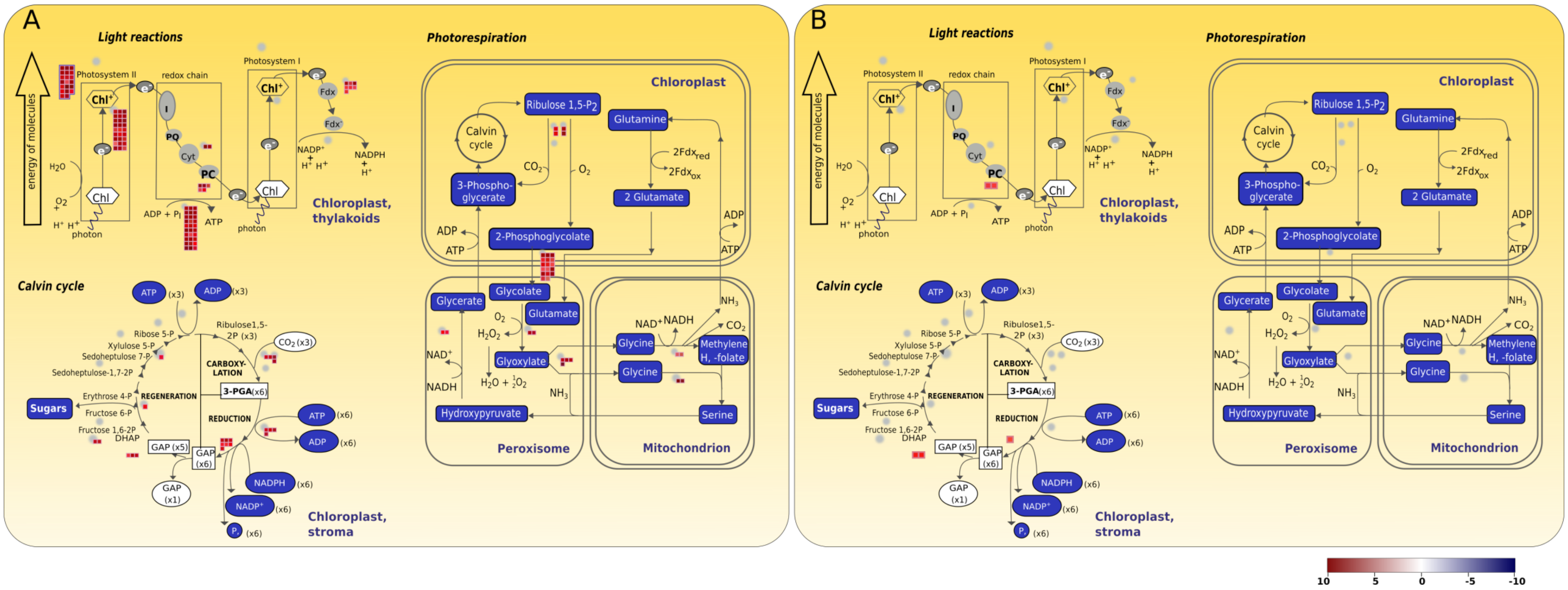


**Supplementary Figure 4:** Pathway analysis of the top 1000 highest expressed genes in leaves and roots. As expected, we found that photosynthesis genes are enriched in the top 1000 highest expressed genes in leaves. The small groups of boxes in each pathway represent genes involved in that process. The color of these boxes ranges from dark red to dark blue representing extremely high expression and low expression, respectively.

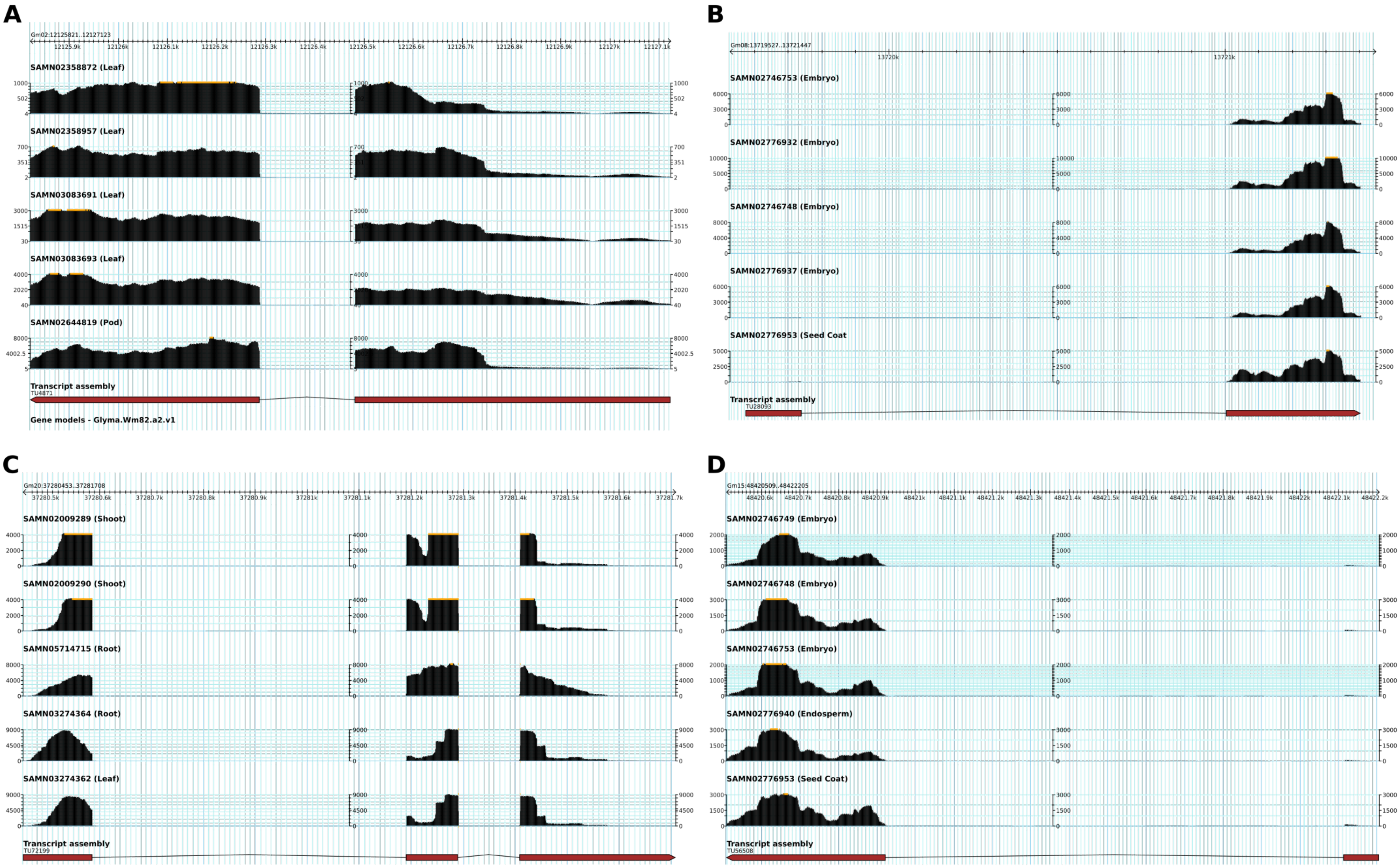


**Supplementary Figure 5:** Wiggle plots showing read coverage of potentially new genes at four unannotated loci on soybean genome. We selected five samples in which these genes had the highest expression levels. A: TU4871, B: TU28093, C: TU72199, D: TU56508.

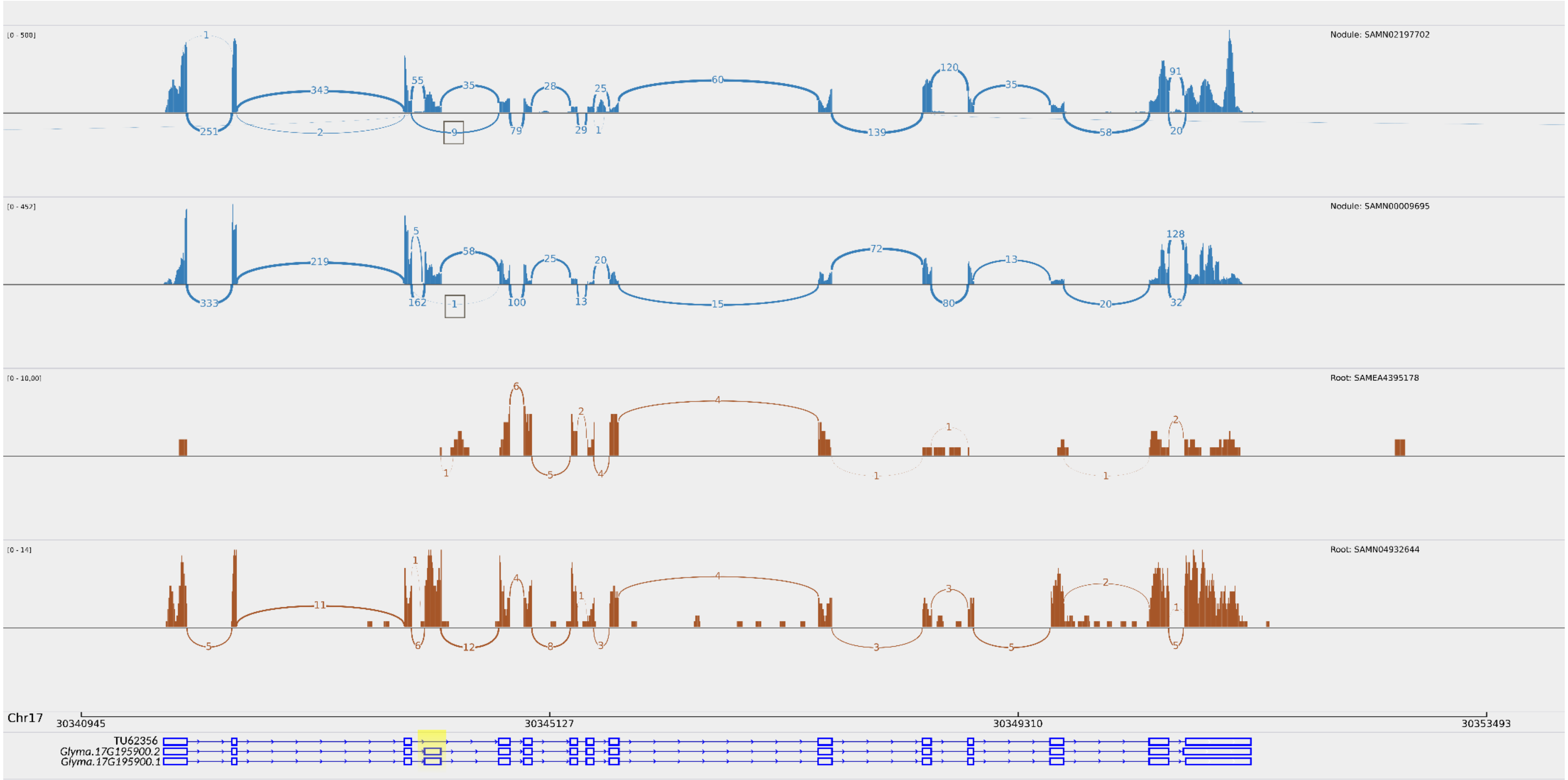


**Supplementary Figure 6:** Sashimi plot of *Glyma.17G195900* (CASEIN KINASE 1-LIKE PROTEIN 4) showing the number of reads supporting splice junctions in two nodule and two root samples. The tracks below the plot represent transcriptional isoforms. Exons within the highlighted region indicate variation in splicing patterns. The top track (TU62356) is the novel isoform, comprising an exon skipping event. From the nodule samples we observed that 10 reads support the skipping of exon 2.

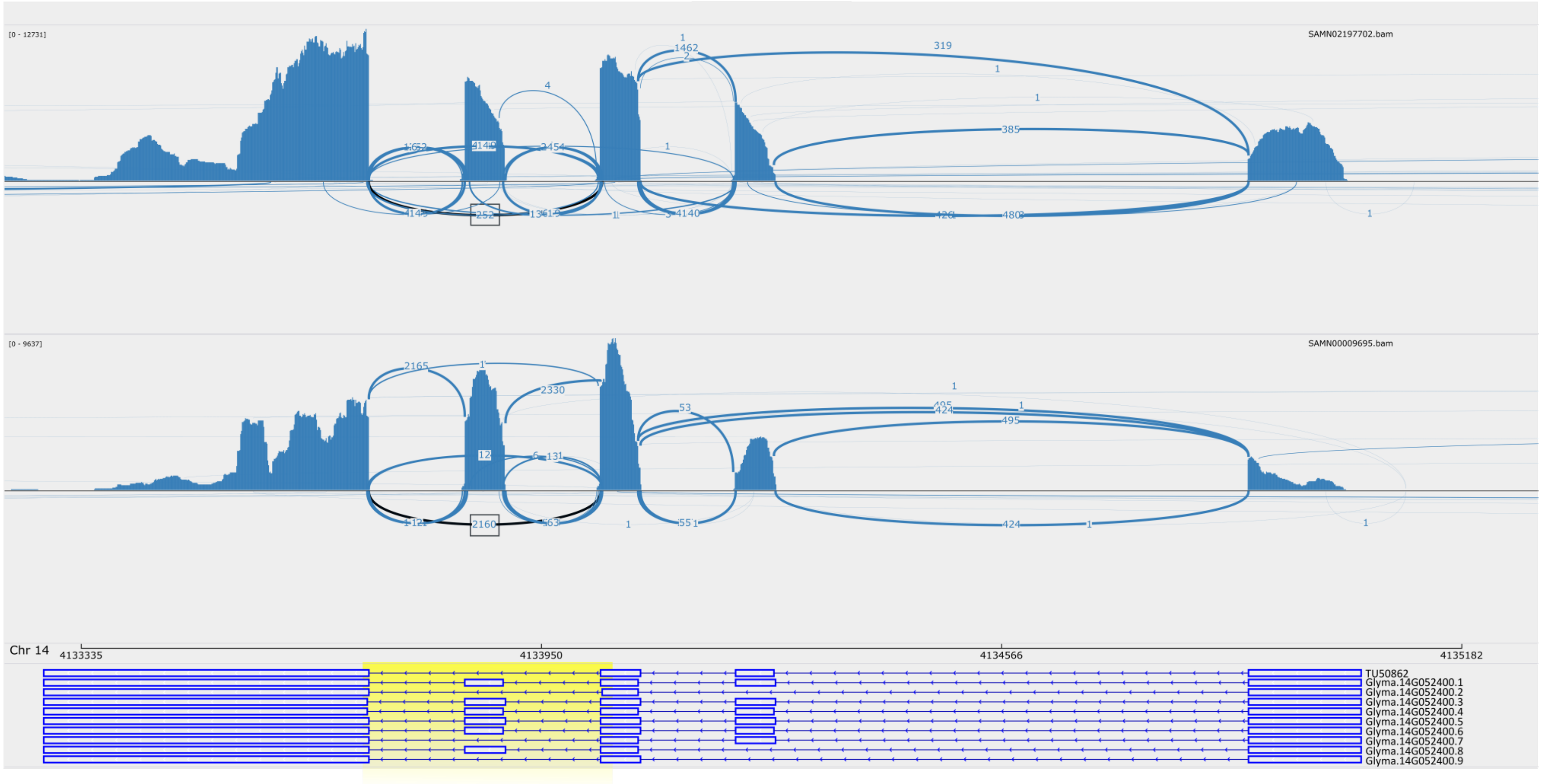


**Supplementary Figure 7:** Sashimi plot of Glyma.14G052400 (Glycine rich protein family) showing the number of reads supporting the splice junctions in two nodule samples. The tracks below the plot represent transcriptional isoforms. The exon within the highlighted region indicate variation in splicing patterns due to the skipping of exon 2 in TU50862. From two nodule samples we observed that 4829 reads have aligned across exons 1 and 3, supporting the skipping of exon 2.

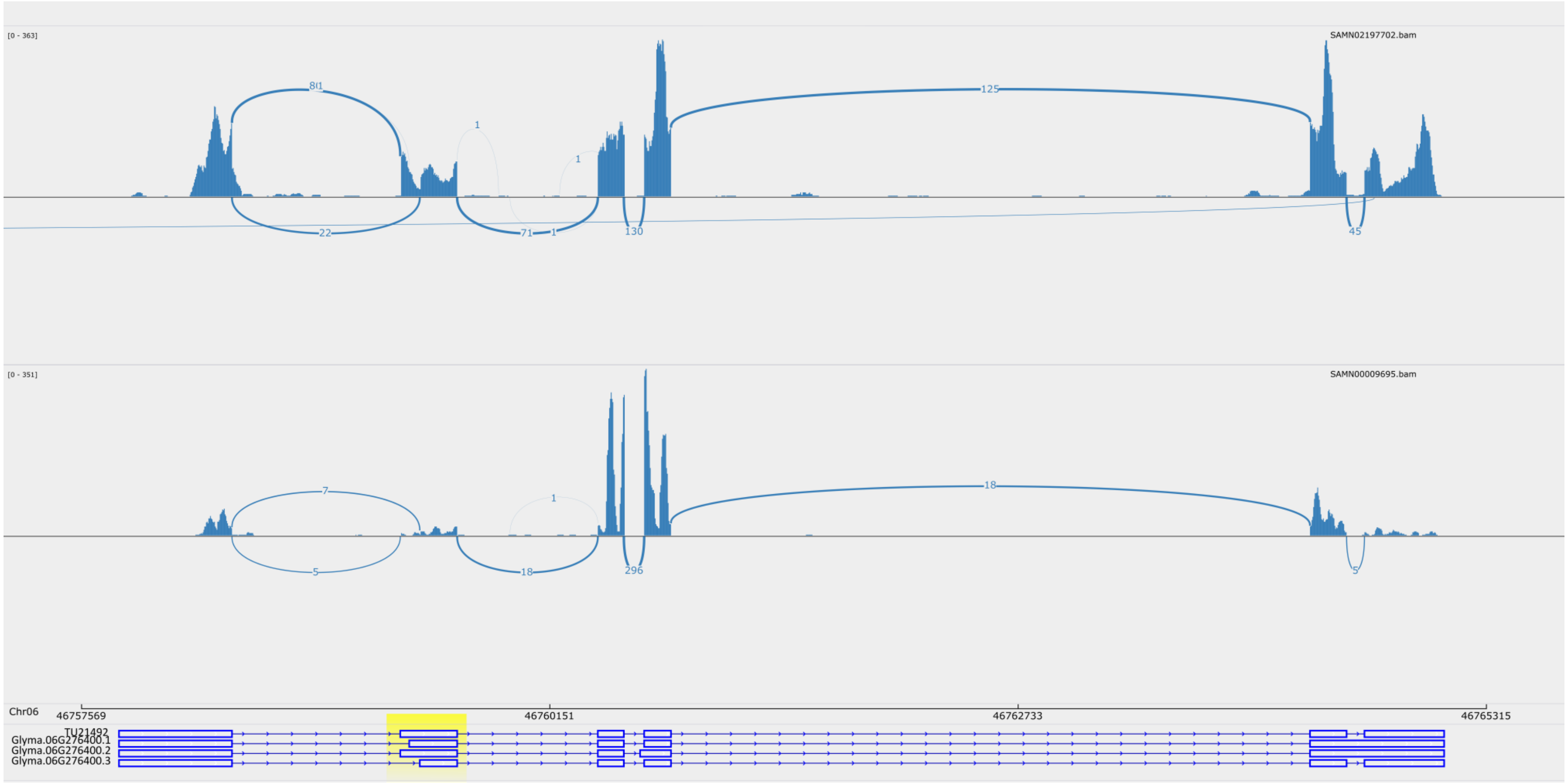


**Supplementary Figure 8:** Sashimi plot of Glyma.06G276400 (Cysteamine dioxygenase/Persulfurase) showing the number of reads supporting the splice junctions in two nodule samples. The tracks below the plot represent transcriptional isoforms. Exons within the highlighted region indicate variation in splicing patterns. The top track (TU21492) is a novel isoform comprising of different lengths of exon 2 as compared to the primary isoform Glyma.06G2764.
